# Differential changes in the effective neural drive following new motor skill acquisition between vastus lateralis and medialis

**DOI:** 10.1101/2025.04.23.650243

**Authors:** Caterina Cosentino, Hélio V. Cabral, Milena A. dos Santos, Elmira Pourreza, J Greig Inglis, Francesco Negro

## Abstract

**Purpose:** To investigate whether short-term learning of a new motor task is mediated by changes in common synaptic inputs to motor neurons within and between synergistic muscles.

**Methods:** Seventeen healthy individuals performed 15 repetitions of a complex force-matching task at 10% of a maximal voluntary contraction. Two trials were selected for analysis, the one with the highest force-target error (pre-learning) and the one with the lowest (post-learning). High-density surface electromyograms recorded from vastus medialis (VM) and vastus lateralis (VL) were decomposed into their constituent motor unit spike trains, with individual motor units being tracked between trials. Motor unit discharge behavior and common synaptic oscillations across the delta, alpha, and beta bands were calculated and compared between pre- and post-learning.

**Results:** Force-target matching improved across trials, accompanied by a significant decrease in the coefficient of variation of the inter-spike interval (*p* < 0.01), while the mean discharge rate remained similar (*p* > 0.85). The area under the curve within delta (*p* < 0.003) and alpha (*p <* 0.004) bands decreased between trials, with no significant changes in the beta band (*p* > 0.05). Notably, reductions in the alpha band correlated significantly with performance improvements in VL (R = 0.81) but not in VM (R = 0.12).

**Conclusion:** The acquisition of a new motor task is mediated by modulations in common synaptic inputs to motor units, leading to improved force control. Our findings further suggest that these changes in common synaptic inputs, particularly in the alpha band, differ between VM and VL.

## Introduction

The acquisition of a new motor skill task is defined as the process by which movements become progressively more accurate through practice and interaction with the environment (Willingham, 1998; Cheung et al., 2020; Doyon et al., 2003). This incremental process begins with a rapid initial phase, during which task repetition within a single session induces specific changes in movement control, ultimately enabling precise execution of the new motor skill (Doyon and Benali, 2005). These early changes in movement control reflect neural plasticity at both cortical and subcortical levels, ensuring the coordinated activation of synergistic muscles required for the intended movement (Dayan and Cohen, 2011; Ungerleider et al., 2002; Ziemann et al., 2004; Stefan et al., 2006; Aizenstein et al., 2004; Hikosaka et al., 2002; Doyon et al., 2002; Floyer-Lea and Matthews, 2004; Cheung et al., 2020). For instance, the early learning phase is dominated by cerebellar activity, with peak activation of the primary motor cortex linked with error correction (Lohse et al. 2014). Thus, research on functional and neural circuitry adaptations at the supraspinal level has significantly advanced our understanding of motor skill learning mechanisms.

Spinal mechanisms underlying voluntary movement generation have also been explored in the context of motor skill learning (Ely et al., 2022; Kinany et al., 2023; Vahdat et al., 2015; Landelle et al., 2021). Compelling evidence suggested that learning a new motor task may modulate presynaptic inhibition by reducing H-reflex excitability, potentially facilitating motor sequence consolidation within spinal circuits (Lungu et al., 2010; Giboin et al., 2020; Perez et al., 2005). Additionally, a recent study investigated how the acquisition of a force-matching task influences the low-frequency oscillations of shared synaptic inputs to alpha motor neurons (Cabral et al., 2024), which are key determinants of force generation and variability (Enoka and Farina, 2021; Hug et al., 2023a; Farina and Negro, 2015, Negro et al., 2009). The findings revealed that learning the force-matching task was associated with reductions in physiological tremor oscillations within the shared synaptic inputs, with these reductions correlating with improvements in performance. However, previous research has primarily focused on individual muscles controlling a single joint. How shared synaptic inputs are transmitted across synergistic muscles and modulated during skill acquisition remains unclear.

Therefore, the aim of this study was to investigate if the short-term learning of a motor task is mediated by changes in the common synaptic inputs to motor neurons of synergistic muscles. We hypothesized that the acquisition of a new skill training task would be similar across synergistic muscles and that the effects of neural drive oscillations on the overall force output will mainly depend on the different anatomical conformation of the muscles, the mechanical constraints, and the functions they perform.

## Methods

### Participants

Seventeen healthy individuals (10 males and 7 females, age: 29 ± 7 years; height: 174 ± 11 cm; weight: 71 ± 19 kg) participated in this study and were deemed suitable for the analyses. All participants were free from neuromuscular pathologies or conditions affecting the lower limb. This study was conducted following the approval of the local ethical committee (number NP5665) and in accordance with the latest Declaration of Helsinki. Before the beginning of experiments, all participants provided written informed consent.

### Experimental protocol

A single experimental session was performed lasting approximately 90 minutes. Participants were seated in an isokinetic dynamometer (HUMACNORM Extremity System, CSMi Solutions, Stoughton, MA) with hip and knee joints positioned at 90° of flexion (0° being fully extended) (**Fig. 1a**). The ankle was secured three centimeters above the malleolus to assess isometric knee extension force.

**Fig. 1.**
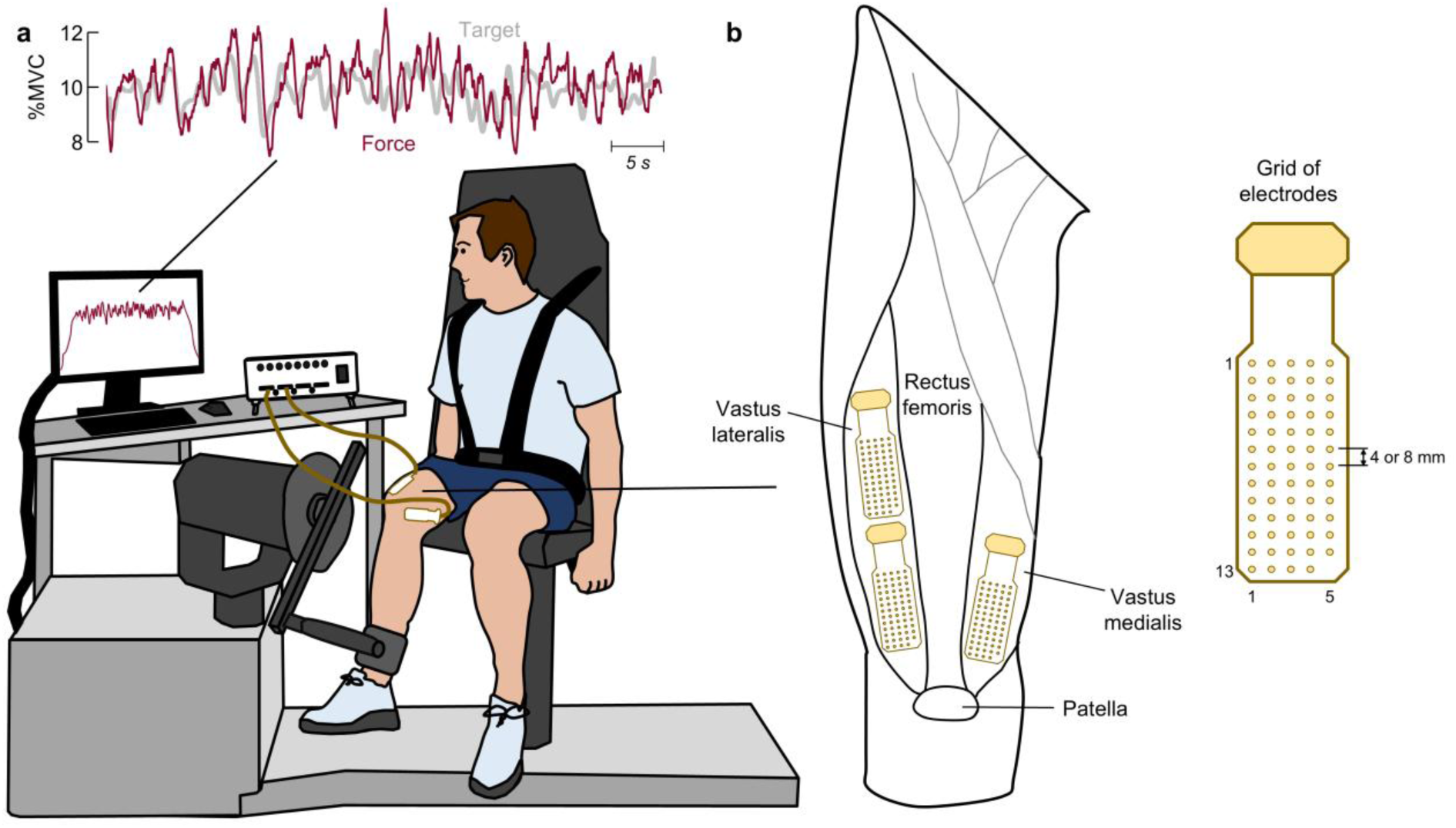
Experimental Setup. (a) Participant’s position in the isokinetic dynamometer for the recording of the isometric knee extension task. Participants were seated on the dynamometer chair with the hip and knee joints positioned at 90° of flexion (0° being fully extended). The learning task consisted of following a randomly generated target signal, which oscillated above and below 10% of maximal voluntary contraction (gray tick line on the top panel). Visual feedback of the force (dark red line on the top panel) was displayed together with the target. (b) Schematic representation of the placement of electrode grids over the vastus lateralis and vastus medialis muscles. Three grids of 64 electrodes arranged in 13 rows x 5 columns were used

First, participants performed three five-second maximal voluntary contractions (MVCs). To ensure that contractions were not influenced by fatigue, a three-minute rest period was provided between each MVC (Inglis and Gabriel, 2020). The maximal force across the three MVCs was used to calculate the force level for the subsequent submaximal contractions (%MVC). Following three minutes of rest, participants performed 15 trials of a complex force-matching task at 10% MVC. The complex force-matching task consisted of a ramp-up phase at a rate of 10% MVC/s, a 30s phase of a stochastic force oscillation, and a ramp-down phase at a rate of 10% MVC/s. During the 30s stochastic force phase, subjects tracked a randomly generated signal with a frequency content below 1.5 Hz (-3 dB low pass frequency), which oscillated above and below 10% MVC (gray tick line on the top of **Fig. 1a**). This protocol was based on prior studies, which have demonstrated clear performance improvements after 15 repetitions of a similar protocol (Knight and Kamen, 2004; Cabral et al., 2024). Visual feedback of the force output was displayed on a monitor along with the target force trace (**Fig. 1a**).

### Data collection

During all 15 trials, the myoelectric signals from vastus medialis (VM) and vastus lateralis (VL) were recorded using high-density surface electromyography (HDsEMG). Specifically, three grids of 64 electrodes arranged in 13 rows x 5 columns (1 mm diameter, 8 mm inter-electrode distance, GR08MM1305, OT Bioelettronica, Turin, Italy) were placed over the distal VM region and the distal and proximal VL regions (**Fig. 1b**). The position of the grids was determined following Barbero et al. (2012). Following a preliminary analysis that revealed a very low number of extracted motor units from the VM muscle (see Results section), for the last 6 participants, three grids of 64 electrodes with smaller inter-electrode distance (4 mm) were applied over the VM, as recent evidence suggests that a larger number of close-spaced electrodes may provide a greater motor units yield (Caillet et al., 2023). Reference electrodes (WS1.1, OT Bioelettronica, Turin, Italy) were positioned on the right patella and the right ankle. Prior to electrode placement, the skin was shaved and mildly abraded (EVERI, Spes Medica, Genova, Italy) and cleaned with water to minimize skin impedance. Double-sided adhesive foam (FOA08MM1305 and FOA04MM1305, OT Bioelettronica, Turin, Italy) was used to attach each matrix to the skin. A thin layer of conductive paste (AC cream, OT Bioelettronica, Turin, Italy) was spread over the electrodes to ensure electrode-skin contact. The myoelectric signals were acquired in a monopolar configuration with a sampling frequency of 2048 Hz using a 16-bit amplifier (10-500 Hz bandwidth; Quattrocento, OT Bioelettronica, Turin, Italy).

### Data Analysis

The analyses were conducted using customized MATLAB scripts (R2023b).

#### Force

The force signals were initially low-pass filtered using a third-order Butterworth filter with a cutoff frequency of 15 Hz. Subsequently, during the 30s force oscillation phase, the root-mean-square error (RMSE), and the cross-correlation between the target profile and the force were computed for all 15 trials. To evaluate improvements in task performance during the force-matching skill acquisition, two out of the 15 trials were selected for each participant. Specifically, the trials selected represented the highest (pre-learning) and lowest (post-learning) RMSE between force output and the target trace (**Fig. 2a**). To assess changes in force steadiness with the force-matching skill acquisition, the coefficient of variation of the force was calculated as the ratio of the standard deviation to the mean value during the 30-s force oscillation phase. Finally, to assess temporal shifts between the feedback and force signals with the force-matching skill, the cross-correlation lag was compared between the pre- and post-learning trials.

**Fig. 2.**
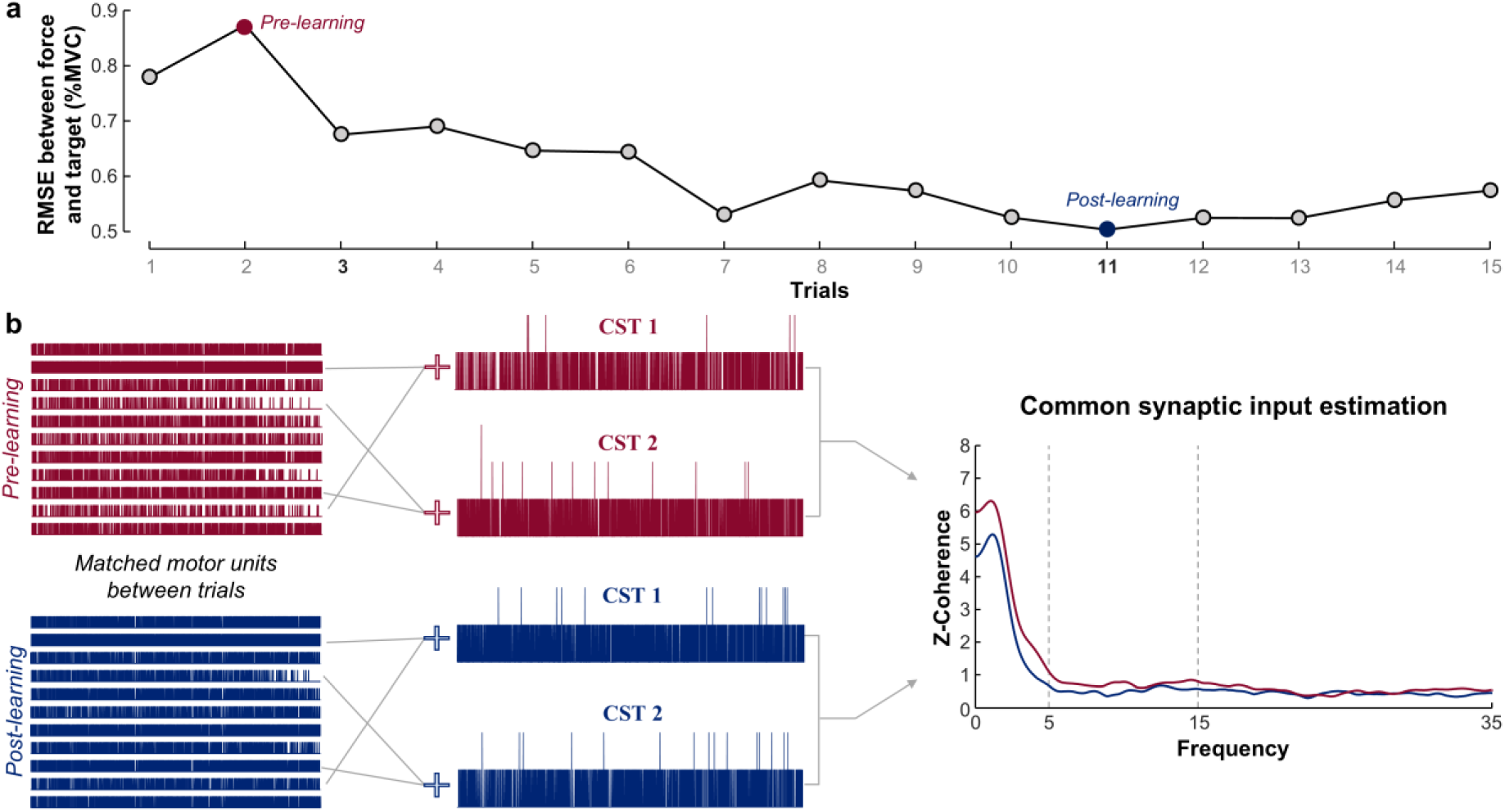
Common synaptic input estimation. (a) Representative case of the root-mean-square error (RMSE) between force and target across the 15 trials of the learning task. Two trials were selected for analysis: the trial with the highest RMSE value (pre-learning; red circle) and the trial with the lowest RMSE value (post-leaning; blue circle). (b) Motor units were decomposed and tracked between the pre-learning (red) and post-learning (blue) trials. Common synaptic oscillations were estimated through coherence analysis between two equally sized cumulative spike trains obtained by summing the discharge times of randomly selected pairs of motor units

#### Motor unit decomposition

HDsEMG signals were initially bandpass filtered using a third order Butterworth filter with cut-off frequencies between 20 and 500 Hz. Subsequently, signals were visually inspected by an expert operator, and noisy channels resulting from electrode contact issues or artifacts were excluded. The signals acquired from each grid during the middle 30s of the target (oscillatory region) were then fully decomposed using a convolutive blind source separation algorithm (Negro et al., 2016). This method allows for the automatic extraction of motor unit spike trains from the HDsEMG signals. The decomposition was performed independently on each electrode grid. Only motor units with a silhouette (SIL) of at least 0.87 were considered for further analyses. The SIL is a measure of accuracy of the decomposition that assesses the separability between identified motor unit spikes and noise (Negro et al., 2016). The identified motor units were visually inspected by the same expert operator. Any missed or misidentified pulses (inter-spike intervals < 20 ms or > 250 ms (Negro et al., 2009)) were manually added or removed based on the changes in the innervation pulse trains and SIL values of motor units. After the manual optimization of the extracted motor units, the occurrence of the same motor unit being recorded by different grids placed on the same muscle (e.g., proximal and distal VL regions) was assessed. In cases duplicated units were identified across grids (i.e., motor units that shared more than 30% of their discharge times), the motor unit with the highest coefficient of variation of the inter-spike interval value was excluded for the subsequent analyses.

#### Motor unit tracking

Following the decomposition, the identified motor units were tracked between the pre- and post-learning trials by reapplying the motor unit separation vectors from the pre-learning to the post-learning trial and vice-versa (Oliveira and Negro, 2021; Cabral et al., 2024). This procedure was performed separately for each electrode grid. For two subjects who did not have at least four units tracked between trials, the signals were further inspected to track the motor units based on the similarity of their shape (i.e., motor unit template matching method) (Martinez-Valdes et al., 2017). For this analysis, only motor units with a cross-correlation between their action potentials greater than 0.8 were considered (Martinez-Valdes et al., 2017). For all tracked motor units between trials, the mean discharge rate and the coefficient of variation of the inter-spike interval were calculated.

#### Common synaptic input within and between muscles

To assess changes in common synaptic inputs to the motor neuron pool following the learning of a new motor skill task, coherence analysis was performed between motor unit spike trains (Castronovo et al., 2015; Negro and Farina, 2012). This analysis measures the linear coupling between two signals in the frequency domain, where zero indicates no coupling and one perfect coupling at a given frequency. Both within-muscle (VM and VL separately) and between-muscle (pooled VM-VL) coherence were performed. Changes in common synaptic oscillations were estimated separately for the delta (1-5 Hz), alpha (5-15 Hz), and beta (15-35 Hz) bands. Specifically, after the discharge times of motor units were converted into binary sequences (i.e., motor unit spike trains), two equally sized cumulative spike trains (CSTs) were obtained by summing the binary sequences of two randomly selected motor units from the matched units (**Fig. 2b**). This process was repeated for up to 100 interactions and the average coherence was calculated for further analysis. As an inclusion criterion, participants needed to have at least four tracked motor units between trials. For all interactions, coherence was calculated between the two detrended CSTs using Welch’s periodogram with a Hanning window of one second and an overlap of 95% (**Fig. 2b**). The averaged coherence values were converted into standard z-scores as suggested by Gallet and Julien (2011). Only z-coherence values greater than the bias, which was determined as the mean z-coherence between 250 and 500 Hz (Castronovo et al., 2015), were considered. To assess changes in the common synaptic inputs across different frequency bandwidths, the area under the curve was calculated for delta, alpha, and beta bands. Subsequently, the ratio of the area under the curve (post/pre) was calculated and subtracted by one. In this way, positive values indicated an increase in coherence within the specific frequency bandwidth, while negative values indicated a decrease.

#### Coupling between force/neural drive and target oscillations

To assess whether there were changes in the linear coupling between the oscillations in force and the target oscillations, as well as between oscillations in the neural drive (i.e., CSTs) and the target oscillations, coherence analysis between these signals was performed. This analysis was carried out similarly to the coherence between motor unit spike trains. However, as force control is strictly associated with low frequencies and the target presented a narrow bandwidth (below 1.5 Hz), only the delta band was considered for this analysis (Negro and Farina, 2012).

### Statistical analysis

Statistical analyses were performed in R (v. 4.3, R Core Team, 2023) using the RStudio environment (v. 2023.03.1). Non-parametric tests were used considering the data was non-normally distributed (Kolmogorov-Smirnov test; *P* < 0.001 for all). The Wilcoxon signed-rank test was performed to compare the force-target RMSE, the force-target cross-correlation values and the force-target cross-correlation lags between trials. The Wilcoxon signed-rank test was also used to compare the coefficient of variation of force between the pre-learning and post-learning trials.

Linear mixed models (LMMs) were employed to compare the mean motor unit discharge rate and the coefficient of variation of the inter-spike interval between pre- and post-learning. LMMs were implemented with the *lmerTest* package (Kuznetsova et al., 2017) using the Kenward-Roger method to approximate the degrees of freedom and estimate *P*-values. For this analysis, random-intercept models were used, considering time (pre-learning and post-learning) as a fixed effect and participant as a random effect. The marginal means with 95% confidence level intervals were estimated using the *emmeans* package (Lenth, 2016). To compare z-coherence values between pre- and post-learning trials, changes in the area under the curve ratio (post/pre) were assessed using the one-sample Wilcoxon test (null hypothesis µ_0_ = 0), separately for each frequency band. To determine whether the changes in the area under the curve of the z-coherence were associated with the number of motor units, Pearson’s correlations were performed between the number of matched motor units and the area under the curve ratios, separately for the delta and alpha bands. For this analysis, the VM and VL results were pooled together. To evaluate changes in the area under the curve ratio of the force-target z-coherence and CST-target z-coherence, the one-sample Wilcoxon test was used.

Separately for each muscle, repeated measures correlations were performed between the force-target RMSE and the area under the curve ratio within the delta and alpha bands to test whether the observed reductions in delta and alpha band coherence (see *Results*) were correlated with performance improvements. These analyses were performed using *rmcorr* package with fixed slopes (Bakdash and Marusich, 2017). For all the variables, statistical significance was set at an alpha of 0.05. All individual data of motor unit discharge times for both the VM and VL acquired in the pre- and post-learning trials are available at https://doi.org/10.6084/m9.figshare.28847492.

## Results

### Changes in force oscillations between pre- and post-learning

To assess improvements in force-matching between pre- and post-learning trials, the RMSE and cross-correlation values between the force output and the target trace were calculated. **Fig. 3a** shows the force traces of a representative participant during the pre-learning (red) and post-learning (blue) trials. It can be visually observed that there is a greater overlap between the force and the target (gray line) in the post-learning compared to the pre-learning, indicating improvements in performance.

**Fig. 3.**
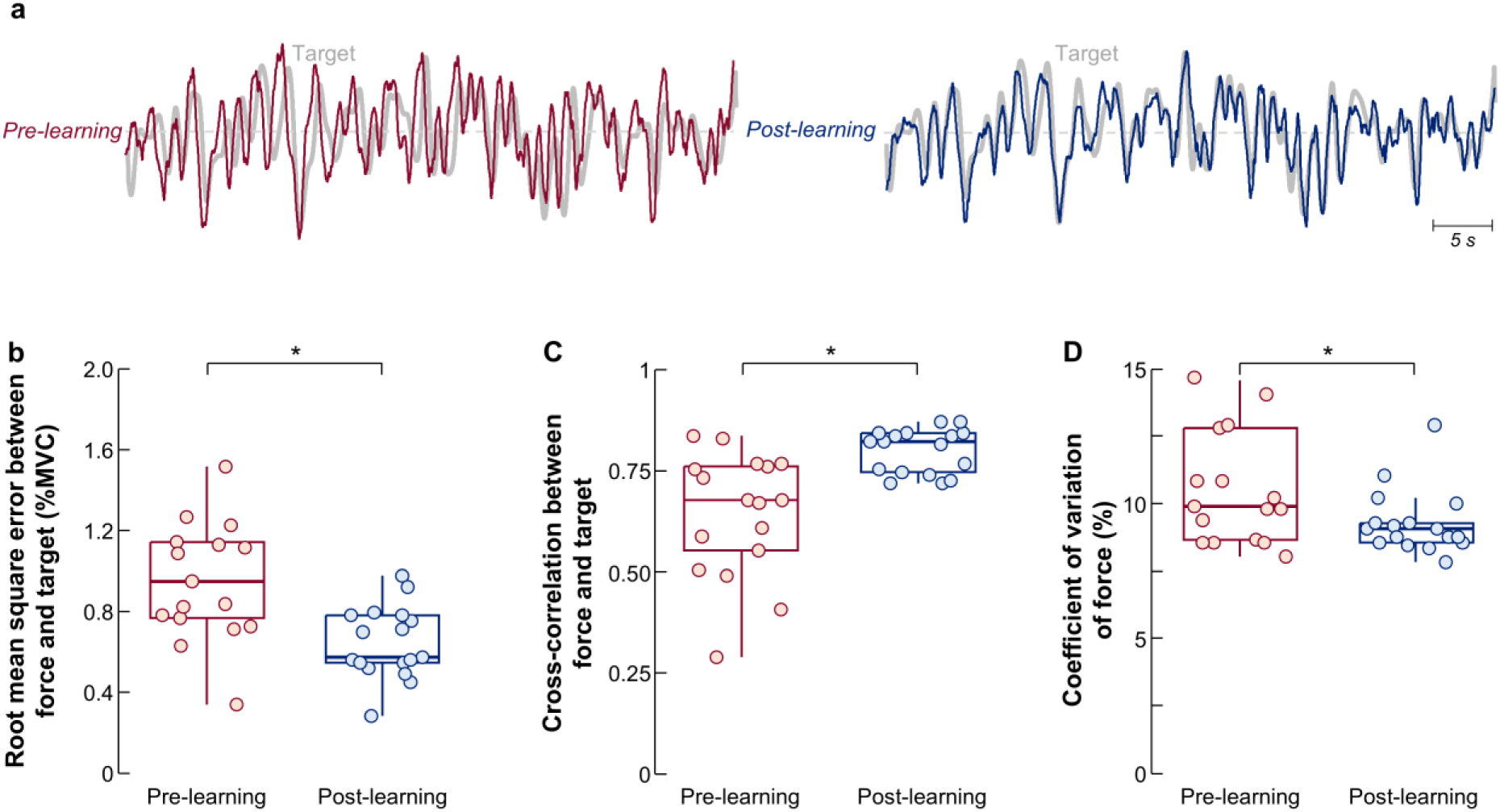
Force-target matching results. (a) Representative case of the force during the pre-learning (red) and post-learning (blue) trials. The target is shown in gray. (b) Group results of the root-mean-square error between the force and the target. (c) Group results of the cross-correlation between the force and the target. (d) Group results of the coefficient of variation of force. Circles identify individual participants. Horizontal traces, boxes, and whiskers denote the median value, interquartile interval, and distribution range. *p < 0.05

These observations were confirmed by the group results. There were statistically significant decreases in RMSE (**Fig. 3b**; *p* < 0.001) and increases in the cross-correlation (**Fig. 3c**; *p* < 0.001) between the force and the target when comparing post-learning with pre-learning trials. However, the cross-correlation lags did not significantly change between the pre- and post-acquisition trials (*p* = 0.47). Moreover, to assess changes in force steadiness, the coefficient of variation of the force was calculated and compared between trials. As shown in **Fig. 3c**, there was a significant decrease in the coefficient of variation of force (*p* = 0.009), which indicates an increase in force steadiness with force-matching skill acquisition.

### Motor unit discharge behavior

A total of 211 motor units were identified from the VM (107 pre-learning; 104 post-learning), with an average (± standard deviation) of 6 ± 3 motor units per participant in the pre-learning (4 ± 2, 3 ± 2 and 3 ± 2 in the three grids over the VM) and 6 ± 4 motor units in the post-learning (5 ± 2, 4 ± 1 and 5 ± 3 in the three grids over the VM). In the VL, 635 motor units were identified (297 pre-learning; 338 post-learning), with an average of 17 ± 8 (7 ± 4 and 10 ± 6 in the two grids over the VL) and 20 ± 10 (9 ± 3 and 13 ± 7 in the two grids over the VL) motor units per participant in the pre- and post-learning trials, respectively. These motor units were subsequently matched between pre- and post-learning trials, leading to an average of 5 ± 3 matched motor units for the VM (3 ± 2, 3 ± 1 and 2 ± 2 in the three grids over the VM) and 15 ± 9 for the VL (4 ± 2 and 10 ± 8 in the two grids over the VL). The average (± standard deviation) SIL values of the matched motor units were 0.88 ± 0.04 and 0.91 ± 0.05 for the VM and VL, respectively.

To assess changes in motor unit discharge behavior following the learning of the force-matching skill task, the mean discharge rate and coefficient of variation of the inter-spike interval of tracked motor units were calculated. For both VM and VL, there were no significant changes in mean discharge rate between pre- and post-learning (VM: from 7.81 [7.24, 8.38] pps to 7.83 [7.26, 8.39] pps, *p* = 0.95; VL: from 8.19 [7.61, 8.77] pps to 8.17 [7.59, 8.75] pps, *p* = 0.87). In contrast, the coefficient of variation of the inter-spike interval significantly decreased between trials for both the VM (from 47.0 [39.4, 54.6] % to 35.6 [28.0, 43.2] %, *p* = 0.01) and VL (from 42.1 [36.9, 47.2] % to 27.4 [22.3, 32.6] %, *p* < 0.001).

### Common synaptic input oscillations

To evaluate alterations in common synaptic oscillations between the pre- and post-learning trials, coherence analysis was conducted between motor unit spike trains, both within and between VM and VL. Specifically, the area under the curve ratio (post/pre) was calculated for the delta (1-5 Hz), alpha (5-15 Hz), and beta (15-35 Hz) bands. Only participants with at least four tracked motor units between trials (see *Methods*) were considered for this analysis; thus, 10, 16 and 17 participants were included for VM, VL and VM-VL coherence, respectively. In both muscles, significant reductions were observed in the area under the curve within the delta (*p* = 0.02 for VM; *p* = 0.003 for VL) and alpha bands (*p* = 0.04 for VM; *p* = 0.004 for VL) but did not significantly change for the beta band (*p* = 0.75 for VM; *p* = 0.06 for VL) (**Fig. 4a and 4b**). Similar results were observed for the VM-VL coherence, with significant decreases in the delta and alpha bands with the force-matching skill acquisition (*p* < 0.05 for both), but not for the beta band (*p* = 0.27; **Fig. 4c**). To assess if observed changes in common synaptic oscillations were associated with the number of included motor units, we performed a Pearson’s correlation analysis between the area under the curve ratio (post/pre) and the number of matched motor units. For both the delta (R = -0.03, *p* = 0.88) and the alpha bands (R = 0.03, *p* = 0.86), the number of motor units was not significantly correlated with changes in the area under the curve.

**Fig. 4.**
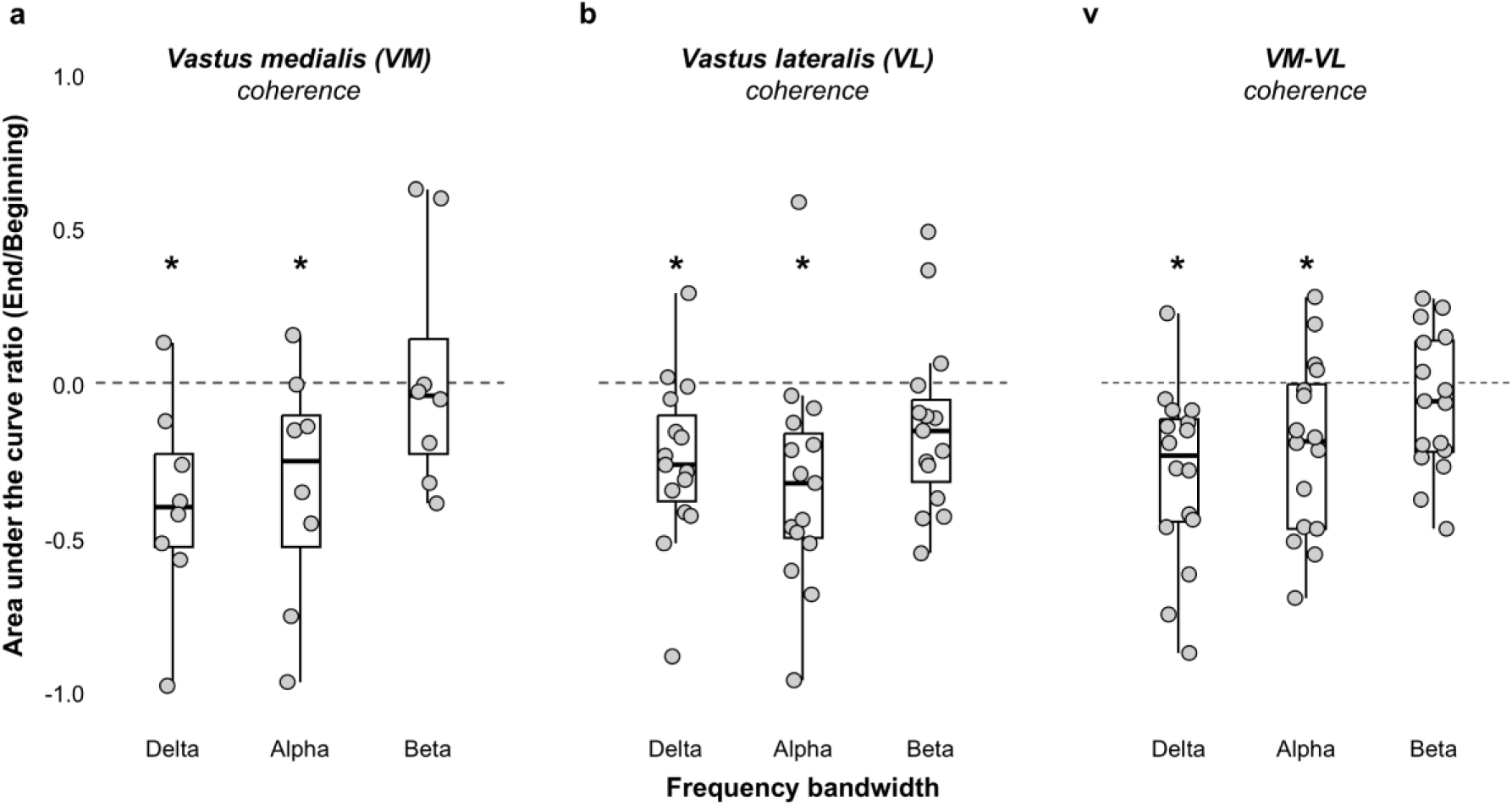
Motor unit coherence results. Group results of the area under the curve ratio of coherence for VM (a), VL (b), and pooled VM-VL muscles (c). Three frequency bandwidths were analyzed: delta (1-5 Hz), alpha (5-15 Hz), and beta (15-35 Hz). Median value, interquartile range, and distribution range are denoted by the horizontal traces, boxes, and whiskers, respectively. Circles represent individual participants

To investigate whether reductions in motor unit z-coherence within the delta and alpha bands were associated with improvements in the force-matching task, we applied a repeated measures correlation. For both muscles, a significant positive correlation was observed between the force-target RMSE and the z-coherence within the delta band (R = 0.54, *p* = 0.03 for the VL; **Fig. 5a**; R = 0.81, *p* = 0.04 for the VM; **Fig. 5c)**. When considering the alpha band, there was a positive correlation between the force-target RMSE and the z-coherence for the VL (R = 0.81; *p* = 0.001; **Fig. 5b**), but not the VM (R = 0.12; *p* = 0.74; **Fig. 5d**).

**Fig. 5.**
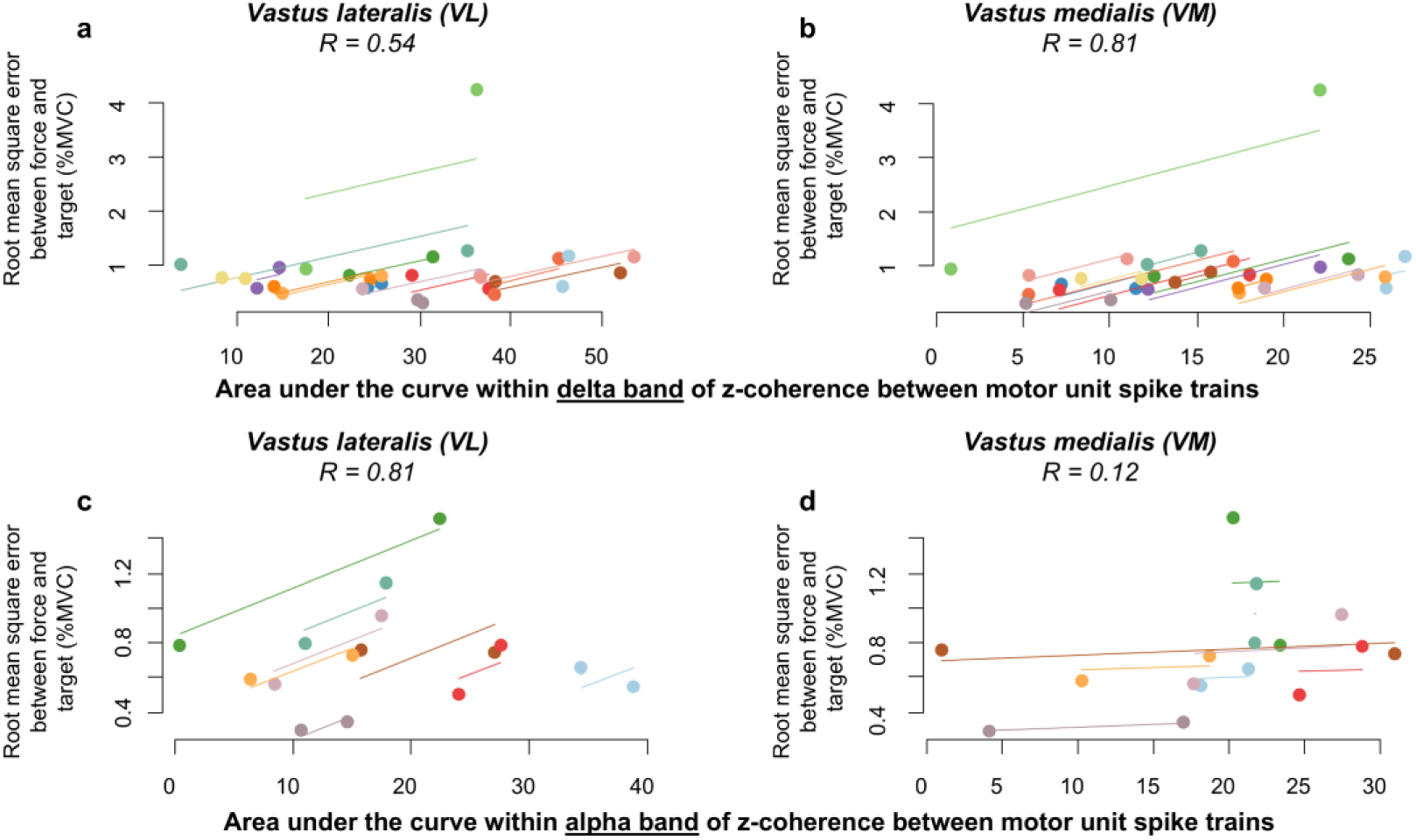
Correlation results. Repeated measures correlation between the force-target root-mean-square error and the area under the curve of motor unit z-coherence within the delta band (a and b), and the alpha band (c and d). The left panel shows the results for the vastus lateralis muscle and the right panel for the vastus medialis muscle

### Linear coupling between force/neural drive and target oscillations

Changes in the linear coupling between the force/neural drive oscillations and the target trace oscillations were investigated using coherence analysis between these signals within the delta band. **Fig. 6** shows the pooled z-coherence between the force and target, and between the neural drive (i.e., CST) and the target for both VM and VL. For both force-target z-coherence (*p* < 0.001; **Fig. 6a**) and VL CST-target z-coherence (*p* = 0.03; **Fig. 6b**), there was a significant increase in the area under the curve between pre- and post-learning trials. In contrast, the z-coherence between the VM CST and target did not significantly change between trials (*p* = 0.64; **Fig. 6c**).

**Fig. 6.**
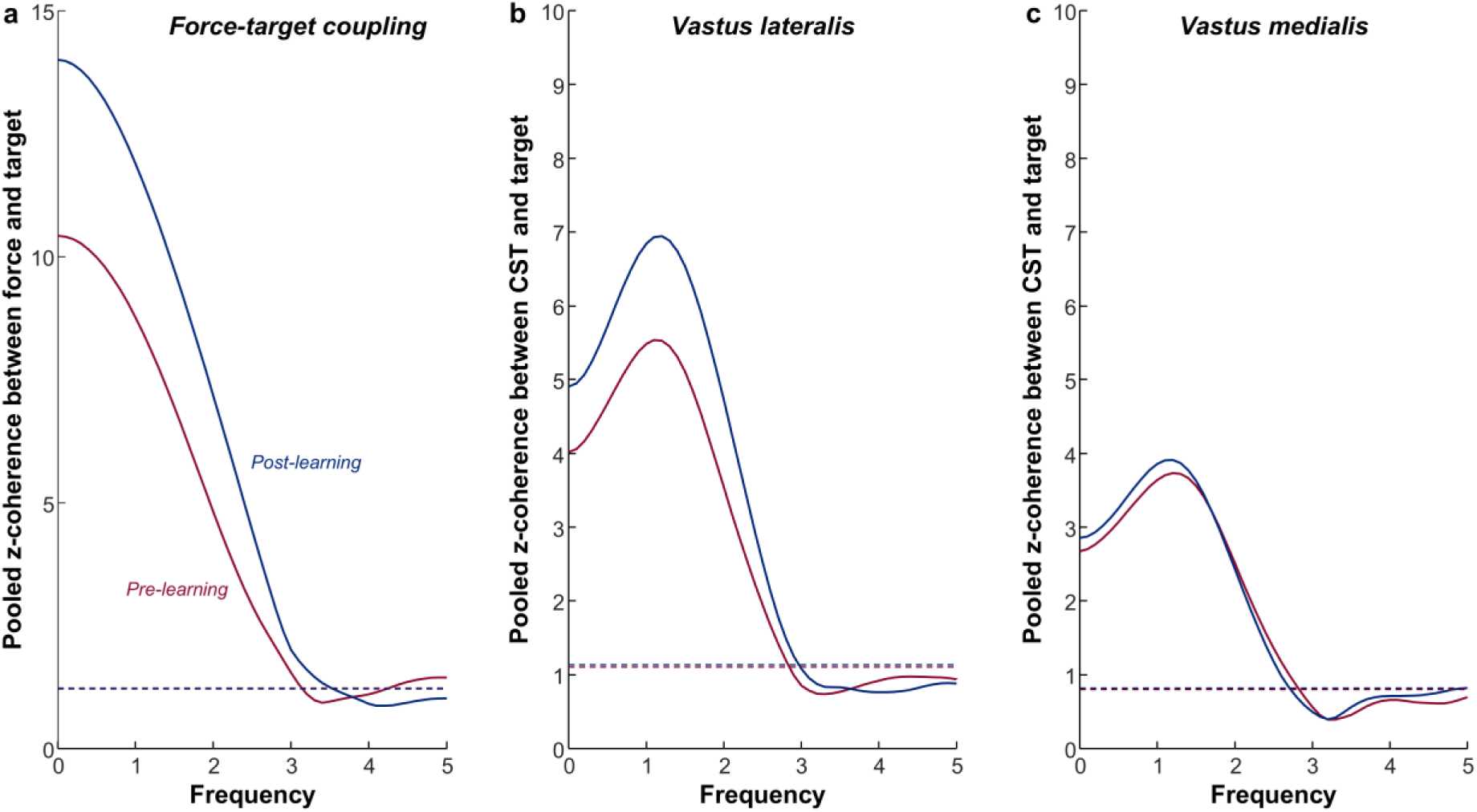
Coherence results between force/neural drive and target oscillations. The pooled z-coherence profiles between force and target (a) and cumulative spike train (CST) and target (b and c) are shown. The CST-target coherence analysis was performed separately for the vastus lateralis (b) and vastus medialis (c) muscles. The red and blue lines denote the pre-learning and post-learning trials, respectively

## Discussion

This study investigated changes in the effective neural drive to synergistic muscles (VM and VL) during the learning of a novel isometric motor task. Our main findings demonstrated that force-matching performance improved with skill acquisition. These improvements were accompanied by a reduction in common synaptic oscillations to spinal motor neurons in the delta and alpha coherence bands, both within and between synergistic muscles. Interestingly, reductions in alpha band oscillations were positively correlated with performance improvements, but only for the VL. As discussed below, these results suggest that changes in neural drive oscillations during short-term motor learning depend on the mechanics and functional roles of synergistic muscles.

Neural inputs transmitted from spinal and supraspinal sources to alpha motor neurons consist of independent inputs linked to the nonlinearities of individual motor neurons, as well as common inputs shared across the entire pool (Negro and Farina, 2012; Castronovo et al., 2018; Enoka and Farina, 2021; Farina and Negro, 2015). Due to the spread of shared inputs, the motor neuron pool functions as a selective linear filter, canceling out input components that are not common to all motor neurons. As a result, the cumulative spike train of action potentials received by the muscle, referred to as the effective neural drive (Farina and Negro, 2015), is primarily determined by the common synaptic inputs (Hug et al., 2023a; Negro et al., 2016; Thompson et al., 2018; Farina et al., 2014; Negro et al., 2009). The neural drive is further lowpass filtered by the contractile properties of active motor units at approximately 12 Hz (Windhorst, 2012). Consequently, low-frequency oscillations in the neural drive (i.e., delta and alpha bands of motor unit coherence) play a determinant role in force control (Farina and Negro, 2015; Hug et al., 2023b). Therefore, we investigated whether learning a new force-matching task is mediated by changes in the effective neural drive to synergistic muscles, by assessing coherent oscillations in motor unit spike trains of VM and VL. Our results revealed that short-term skill acquisition was followed by reductions in alpha band coherence both within and between vastii motor units (**Fig. 4**). Considering alpha band (physiological tremor) oscillations are considered an involuntary noise component of the common synaptic inputs that negatively influence force accuracy (McAuley and Marsden, 2000; Yavuz et al., 2015; Watanabe et al., 2018; Erimaki et al., 1999), their reduction during skill acquisition likely represents a neural mechanism to enhance force precision (Laine et al., 2014; Henderson et al., 2022; Williams et al., 2010; Koželj and Baker, 2014). These findings support previous studies that demonstrated a strong link between reduced physiological tremor oscillations and improved force control (Novak and Newell, 2017; Christakos et al., 2006; Williams et al., 2010; Koželj & Baker, 2014; Laine et al., 2016). Although the precise mechanisms underlying alpha band reduction during short-term learning remain unclear, spinal interneurons phase-cancelling descending alpha oscillations have been proposed as a plausible mechanism (Cabral et al., 2024).

Interestingly, nonetheless similar results in coherence were found when pooling VM and VL motor units (see **Fig. 4**), reductions in alpha band coherence were positively correlated with performance improvements only for VL (**Fig. 5c**), but not VM (**Fig. 5d**). Given that VM and VL are synergistic muscles that share most of their synaptic inputs (∼75-95%; Laine et al., 2015), this finding was unexpected. However, architectural differences between these muscles may explain this discrepancy. Although both VM and VL are part of the same muscle group (i.e., quadriceps), their muscle architecture differs significantly (de Souza et al., 2018; Lieber and Fridén, 2000; Ward et al., 2009; Lefebvre et al., 2006). The VL has a larger cross-sectional area, with muscle fibers arranged more transversely from the distal to the proximal region (Blazevich et al., 2006). These anatomical features provide the VL with a mechanical advantage for force production, while the VM primarily contributes to knee joint stabilization and force transmission to the tendon (De Souza et al., 2018; Lin et al., 2004; Lefebvre et al., 2006; Farahmand et al., 1998; Lieber et al., 2000). Thus, the positive correlation between alpha band reductions and performance improvements only in VL suggests a mechanical advantage to produce force facilitates more efficient transduction of neural inputs into force. This interpretation was further supported by the CST-target coupling analysis, which revealed a significant increase in coupling between pre- and post-learning trials for the VL (**Fig. 6b**), but not for the VM (**Fig. 6c**). Collectively, our results suggest that while synergistic muscles such as the vastii share a significant portion of their synaptic inputs (Laine et al., 2015), their motor output is influenced by their distinct contractile properties and force-generating capacity. These findings highlight the importance of considering muscle-specific functional roles when investigating neural drive adaptations during motor learning.

Our findings also revealed a decrease in common synaptic oscillations within the delta band following skill learning, which plays a critical role in optimal force production and task execution accuracy (Negro et al., 2009). Notably, these reductions in delta band oscillations were positively correlated with improvements in task execution for both VM and VL (**Fig. 5a and 5b**). A plausible explanation for the delta band reductions observed in this study relates to the motor strategy adopted by participants during the learning of the force-matching isometric knee extension task. In both pre- and post-learning trials, participants followed similarly the target in time, as confirmed by the absence of significant differences in force-target cross-correlation lags (see *Results*). Thus, improvements in force-matching accuracy were achieved by decreases in force amplitude oscillations, which was confirmed by significant decreases in the coefficient of variation of force. This reduction in force variability, together with the increased coupling between force and feedback across trials, explains the delta band coherence reductions.

A final consideration concerns alterations in motor unit discharge behavior during task learning. Unlike previous studies (Knight and Kamen, 2004; Patten and Kamen, 2000; Cabral, et al., 2024), no significant changes in mean discharge rate between pre- and post-learning trials were observed. This suggests that mean discharge rate alone may not explain the observed changes in common synaptic oscillations. Instead, the reduction in motor unit discharge variability between trials, quantified by the coefficient of variation of the inter-spike interval, contributed to the observed changes in common synaptic oscillations. This aligns with previous findings demonstrating a correlation between reductions in common synaptic oscillations and decreases in the coefficient of variation of the inter-spike interval (Heckman et al., 2009). Consistent with these findings, numerous studies have suggested that improvements in force steadiness are primarily driven by modulation in the common drive to motor neurons, rather than changes in motor unit discharge rate (Semmler et al., 2006; Dideriksen et al., 2012; Bilodeau, 2000; Tracy et al., 2004; Marmon et al., 2011; Farina and Negro, 2015)

In conclusion, this study demonstrated that, similar to individual muscles, the acquisition of a new motor task is mediated by modulations in the common synaptic inputs to motor units, leading to improved force control and short-term motor learning. Notably, our findings suggest that these changes in common synaptic oscillations are influenced by the individual neural and mechanical properties of synergistic muscles, highlighting the importance of muscle-specific adaptations in motor learning.

## Authors’ Contributions

All the authors contributed substantially to the manuscript. Conception and design of the experiments (CC, HVC, FN), collection of data (CC, MAS, EP), analysis of data (CC, HVC, MAS, EP, JGI, FN), interpretation of data (CC, HVC, FN), drafting the article and revising it critically for important intellectual content (CC, HVC, JGI, FN), and final approval of the version to be published (CC, HVC, MAS, EP, JGI, FN). All authors have read and approved the final version of the manuscript and agree with the order of presentation of the authors.

## Statements and Declarations

None conflicts of interest, financial or otherwise, are declared by the authors.

## Acknowledgments

This study was funded by the European Research Council Consolidator Grant INcEPTION contract no. 101045605.

## Abbreviation list

CST: Cumulative spike train
HDsEMG: High-density surface electromyography
LMM: Linear mixed model
MVC: Maximal voluntary contraction
RMSE: Root-mean-square error
SIL: Silhouette
VL: Vastus lateralis
VM: Vastus medialis

## Notes

https://doi.org/10.6084/m9.figshare.28847492

### Competing Interest Statement

The authors have declared no competing interest.

